# Biochemical engineering of 5hmdC-DNA using a Tet3 double-mutant

**DOI:** 10.1101/2025.06.11.658802

**Authors:** Hanife Sahin, Shariful Islam, A Hyeon Lee, Irene Ponzo, Andreas Reichl, Thomas Carell, Pascal Giehr

## Abstract

5-Hydroxymethyl-2’-deoxycytidine (5hmdC) is an important epigenetic marker involved in gene regulation and DNA demethylation. It has potential use as a biomarker for cancer and other diseases due to its significant depletion in various cancers and disease models. This research aimed to develop a reliable and efficient method for generating 5hmdC-containing DNA, addressing limitations in existing techniques. We created a Tet3 stalling mutant that converts 5-methyl-2′-deoxycytidine (5mdC) into a mixture of 5hmdC and 5-formyl-2′-deoxycytidine (5fdC), followed by a reduction step to convert 5fdC to 5hmdC, ensuring a pure 5hmdC state within the CpG context. This method can convert any PCR product, synthetic oligos, and entire genomes into 5hmdC-modified DNA. The principal results demonstrate high specificity and efficiency, providing a robust tool for epigenetic research, cancer diagnostics, and protein binding assays. Additionally, our technique offers 5hmdC-DNA for functional studies and as standards for diagnostic assays.

## INTRODUCTION

In mammals, the stepwise oxidation of 5-methyl-2′-deoxycytidine (5mdC) by Ten-eleven translocation dioxygenases (TETs) leads to the formation of 5-hydroxymethyl-2′-deoxycytidine (5hmdC), 5-formyl-2′-deoxycytidine (5fdC), and eventually 5-carboxy-2′-deoxycytidine (5-cadC).^1–3^ These oxidized cytosine forms (oxCs) are essential intermediates during DNA demethylation, and significant amounts of oxCs have been found in several cell types.^4–6^ Particularly, 5hmdC, now considered the sixth base of the genome, is present in considerable quantities in neuronal cells of the central nervous system and embryonic stem cells.^4^ 5hmdC has been implicated in selective protein recruitment, impacting chromatin structure and thus gene regulation.^7^ Furthermore, 5hmdC is being investigated as a malignancy biomarker due to its depletion in many cancer cell lines.^8^

Standardized modified oligonucleotides are of great importance for studying the biological function of a modification or establishing a particular DNA modification as a biomarker. Modified DNA is used in several biological assays and applications. It is widely used in DNA-protein interaction studies, such as EMSA or pull-down assays, to identify reader proteins of DNA modifications.^9,10^ Furthermore, modified oligonucleotides serve as positive control and calibration standards for Dot blot assays, qPCR, and, most importantly, next-generation sequencing-based detection methods such as BS, oxBS-seq, or third-generation sequencing-based detection methods such as nanopore sequencing.^11–13^ Particularly in NGS-based techniques, which rely on altered base pairing during PCR (BS- and oxBS-seq), following chemical treatment, modified oligonucleotides inform about the conversion rate of the applied chemistry. Lastly, modified DNA has also been used to determine the impact of DNA modification and nucleosome positioning, as well as the binding of proteins to chromatin containing DNA modifications.^14–16^ However, the current limitations of introducing 5hmdC into DNA do not allow for the capture of its native occurrence and the complexity of the underlying DNA sequence.

Currently, there are two ways of generating 5hmdC-containing DNA. Using solid-phase synthesis, commercially available 5hmdC phosphoramidites can be incorporated into single-stranded oligonucleotides, which enables precise control over DNA sequence and modification site.^17^ However, synthesizing oligonucleotides containing 5hmdC is expensive, limited in length, and only a very few 5hmdC positions can be introduced. For longer DNA fragments, PCR amplification with 5hmdCTP is used by substituting the standard dCTP in the reaction.^18^ One complication is that 5hmdC is incorporated at random positions, and not all polymerases accept 5hmdC as a substrate. Overall, these approaches fail to produce oligonucleotides that capture the natural occurrence of 5hmdC in CpG motifs (PCR) and the underlying sequence complexity of the mammalian genome (solid-phase synthesis).

We rationalized that the optimal way of generating 5hmdC-containing DNA would be the enzymatic introduction of 5mdC in a CpG context using M.SssI, followed by the oxidation of 5mdC to 5hmdC. However, selective and complete conversion of 5mdC to only one of the intermediate products, 5hmdC or 5fdC, remains a significant challenge and has not yet been achieved.

Here, we use a truncated mouse Tet3 stalling mutant containing two amino acid substitutions (T940A and Y1567F) to oxidize 5mdC introduced at CpG sites by M.SssI. The double-mutant T940A/Y1567F Tet3 (DM-Tet3) exhibits reduced oxidation efficiency, and 5mdC is oxidized to 5hmdC and 5fdC, with only traces of 5cadC, under controlled or selective reaction conditions. Subsequently, we apply NaBH_4_ reduction, as previously described by Booth et al., to reduce 5fdC to 5hmdC.^19^ This results in DNA containing almost exclusively 5hmdC with a purity of up to 97.7%. This approach allows the conversion of methylated synthetic oligomers, PCR products, and even genomic DNA into a 5hmdC state. By using distinct methyltransferases, 5hmdC can be introduced in various sequence contexts.

This approach enhances the efficiency and specificity of generating 5hmdC-containing DNA, capturing the natural occurrence of 5hmdC, and the sequence complexity of the mammalian genome. Therefore, these probes are best suited for the application of diagnostic assays and for exploring the complex roles of 5hmdC in various biological contexts.

## RESULTS

There is a demand for 5hmdC-containing DNA in science and clinical diagnostics. However, current approaches for preparing such DNA have limitations. We reasoned that the best way to generate 5hmdC-containing DNA would be the enzymatic oxidation of 5mdC, which can easily be introduced by DNA methyltransferases, to 5hmdC. However, selective and complete conversion of 5mdC to only one of the intermediate products, 5hmdC or 5fdC, remains a significant challenge and has not yet been achieved.

In mammalian Tets, a critical active site scaffold consisting of the highly conserved residues-Y1902, which is involved in base-stacking interactions with the pyrimidine base of the inserted target 5mdC,^20^ and T1372 was reported to be crucial for the efficient sequential oxidation of 5mdC to the higher-order oxidation products 5fdC and 5cadC by human Tet2.^21^ Molecular dynamics (MD) simulations have shown that mutations of these two residues alter and reconfigure interactions at the active site, thereby preventing higher-order oxidation of 5mdC to 5fdC and 5cadC. *In vitro* assays further confirmed that these Tet2 mutants either effectively stall at 5hmdC (T1372E) albeit with considerable amounts of 5mdC remaining unreacted (>50% 5mdC) or mutants that permit the formation of significant quantities of 5hmdC and 5fdC (T1372A and T1372A/Y1902F) and to a much lesser extent of 5cadC (T1372A, ∼10% 5cadC; T1372A/Y1902F, no cadC) with little 5mdC remaining.^21^

Since both residues are conserved across mouse and human TET enzymes, we concluded that these mutations should also affect the oxidation efficiency of mouse Tet3. Thus, we introduced mutations (T1372A, T1372E and T1372A/Y1902F) in our recently reported truncated mouse Tet3 variant, hpTet3 (high-performance Tet3), which correspond to T940A, T940E and T940A/Y1567F in hpTet3 (Figure 1A). The N-terminally Strep(II)-tagged mutant proteins, single mutants (SM) T940A and T940E, and double-mutant (DM) T940A/Y1567F Tet3, were overexpressed in *E. coli* and purified in a two-step purification protocol as previously reported for hpTet3.^22^

**Figure 1:**
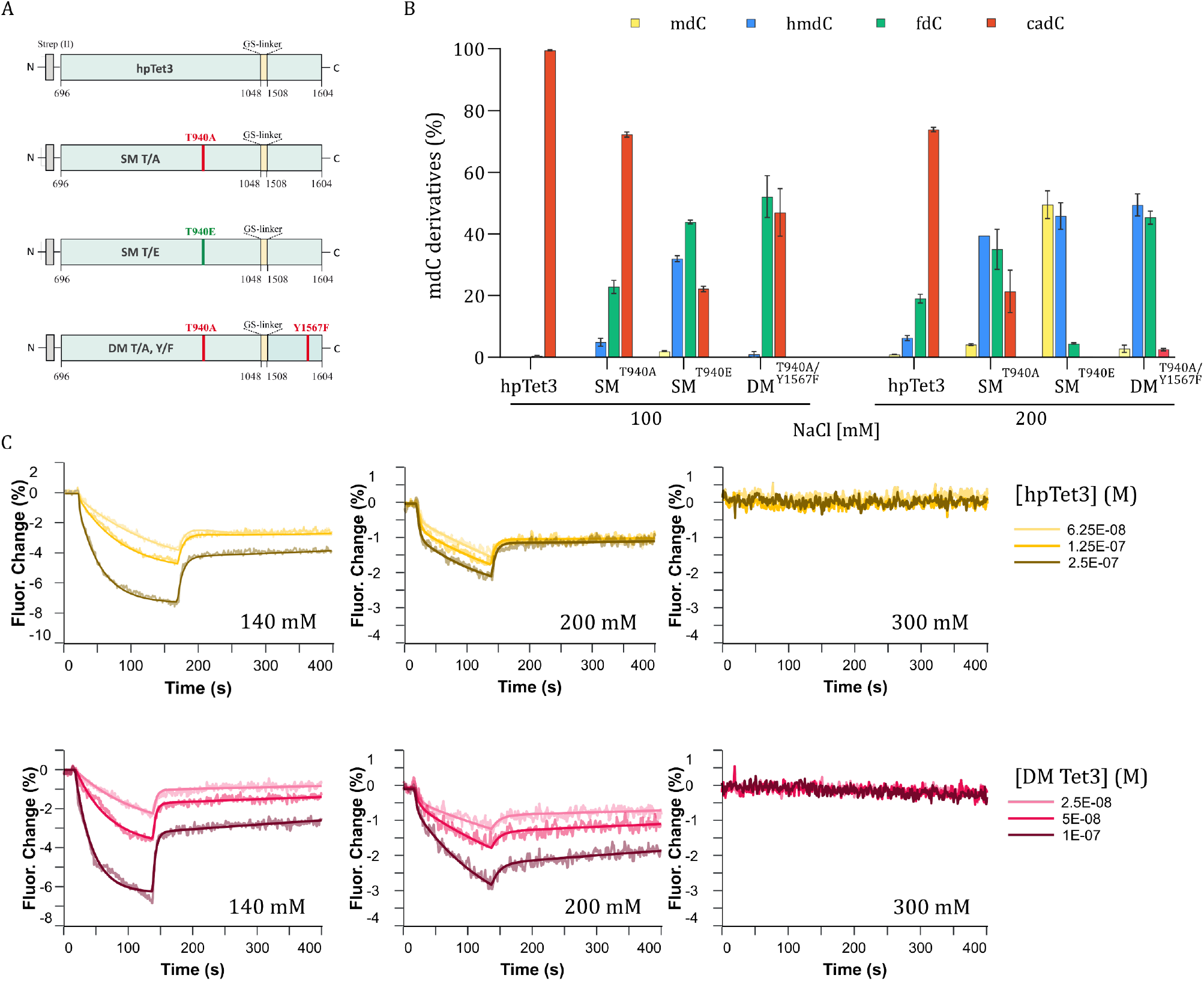
Double-Mutant Structure and Initial Characterization. (A) Schematic representation of the truncated hpTet3, single mutants (SM) carrying the T940A (T/A) or T940E (T/E) mutations, and double-mutant (DM) carrying the T940A (T/A) and Y1567F (Y/F) mutations. (B) UHPLC-QQQ-MS/MS result of: Left - oxidation of mdC by Tet enzyme variants in a buffer containing 100 mM NaCl; Right - oxidation of 5mdC by Tet3 enzyme variants in a buffer containing 200 mM NaCl. Oxidation reactions were performed using 1 µg of human gDNA isolated from HEK293T and 4 µM of the respective enzyme. Values represent mean ± SD, n =4. (C) Binding kinetics of hpTet3 (yellow traces) and DM-Tet3 (pink traces) to 5mdC-containing DNA were performed at different salt concentrations. Top: hpTet3 kinetics measured in buffer containing NaCl 140mM, 200mM, 300mM. Bottom: DM-Tet3 kinetics measured in buffer containing NaCl 140mM, 200mM, 300mM. The enzyme was injected on the surface at the specified concentrations and measured at 25°C.

Next, their oxidation capabilities to stall oxidation at particular oxidative states, preferably 5hmdC, were evaluated *in vitro* at the genomic level using human genomic DNA from HEK293T to account for a wide variety of sequence contexts (…NpNpCpGpNpN…). Notably, oxidation reactions were performed under buffer conditions described for hpTet3, but with 100 mM NaCl as a minor modification, as done by Liu *et. al*.^21^. To that end, 1 µg of HEK293T gDNA was treated with 4 µM of the respective recombinant mutant enzyme for 1 h at 37 °C (for further details, see supporting information). The DNA was purified and digested to the single-nucleoside level and subjected to UHPLC triple quadrupole mass spectrometry (UHPLC-QQQ-MS/MS) using isotopically labelled standards for exact quantification by isotope dilution mass spectrometry.

UHPLC-QQQ-MS/MS analysis of the nucleoside mixture of the SM T940A Tet3 and DM T940A/Y1567F Tet3 revealed that nearly all 5mdCs in the genome were converted to primarily 5fdC and 5cadC, with only ∼5% and ∼1% 5hmdC remaining, respectively. The major oxidation product of SM T940E Tet3 was 5fdC (Figure 1B, left). Overall, all mutants displayed high catalytic activity, permitting substantial formation of the higher-order oxidation products 5fdC and 5cadC. However, we observed an apparent reduction in catalytic activity in all mutants compared to hpTet3, which produced almost exclusively 5cadC (>99%) under the same reaction conditions.

Previously, in an effort to optimize the buffer composition and thereby increase the oxidation efficiency of hpTet3, we determined the optimal ionic strength for hpTet3. Here, we observed that high salt concentrations largely inhibited the oxidation to 5cadC (SI Figure S1). hpTet3 exhibited enhanced activity at lower NaCl concentrations, while substantially reduced activity at higher NaCl concentrations (>100 mM) was observed. We determined the apparent salt optimum for catalysis by hpTet3 with all 5mdCs being converted to 5cadC (>99.9%) to be 50 mM NaCl (SI Figure 1). Importantly, binding kinetics experiments of hpTet3 to 5mdC containing dsDNA revealed that the affinity of the enzyme to DNA is negatively affected by the presence of higher salt concentrations (>200 mM NaCl), where binding to the DNA substrate is progressively disrupted by an increase in salt concentration (Figure 1C, see SI Figure S2 and S3 for experimental details and Table S1 for an overview of the kinetic rates). Thus, in a next step, to fine-tune the catalytic activity and to steer the reaction into preferential formation of 5hmdC, the influence of various salt concentrations (ranging from 50 – 350 mM NaCl) on the formation of the higher-order oxidation products 5hmdC and 5fdC was studied for the mutant Tet enzymes. UHPLC-QQQ-MS/MS revealed that the catalytic activity of all mutants is markedly decreased at elevated salt concentrations, with significant activity loss at salt concentrations higher than 250 mM (SI Figure S4, S5, and S6). Therefore, we focused on 200 mM NaCl as a buffer concentration.

With a NaCl concentration of 200 mM, MS data showed that the double-mutant (DM) T940A/Y1567F Tet3 selectively oxidizes methylated-CpG sites in the HEK292T genome predominantly to 5hmdC and 5fdC. More precisely, at 200 mM NaCl, DM T940A/Y1567F Tet3 robustly produced ∼95% 5hmdC and 5fdC, with only small amounts of 5cadC (2.5%) formed and 5mdC (2.8%) remaining (Figure 1B). This suggests that the ionic strength applied in the reaction modulates the distribution of oxidation products. A noticeable accumulation of 5hmdC and 5fdC was also seen in the SM T940E at 125 mM NaCl (SI Figure 7). However, the sum of 5hmdC and 5fdC was only ∼85%.

To further characterize this mutant and to optimize the formation of 5hmdC, oxidation reactions were repeated with increasing amounts of DM T940A/Y1567F Tet3, ranging from 0.5–8 µM, to determine the maximum extent of DM-Tet3s activity at 200 mM NaCl. UHPLC-QQQ-MS/MS analysis revealed that a distribution favoring the production of ∼95% 5hmdC and 5fdC can be achieved at DM T940A/Y1567F Tet3 enzyme concentrations > 3 µM, while levels of 5cadC and 5mdC remain low (Figure 2A).

**Figure 2:**
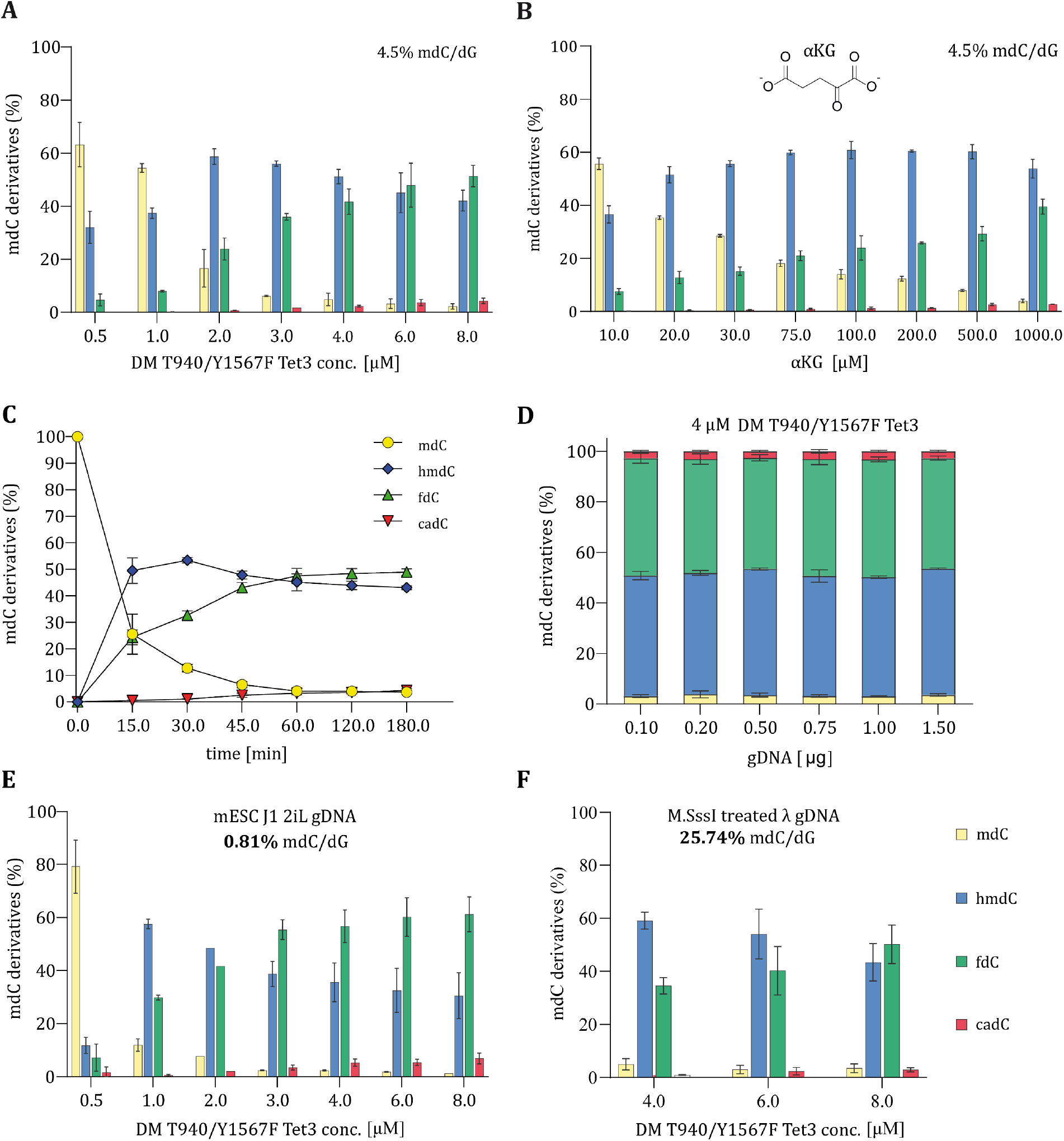
Optimization of the DM-Tet3 reaction conditions. All reactions were conducted using 200 mM NaCl as a buffer condition. UHPLC-QQQ-MS/MS results of (A) optimization of the DM-Tet3 concentration using 1 µg of human gDNA isolated from HEK293T. n=2, (B) evaluation of distinct Alpha-ketoglutarate (αKG) concentration using 1 µg of human gDNA isolated from HEK293T. n=2, (C) distinct incubation times using HEK293T gDNA. n=4, (D) variable genomic DNA (gDNA) concentration derived from HEK293T. n=3, (E) optimization of the DM-Tet3 concentration using 1 µg of gDNA from J1 mouse embryonic stem cells (mESC J1) cultivated with 2iL, and (F) optimization of the DM-Tet3 concentration using 1 µg of M.SssI-methylated DNA from lambda phage extracted from *dam*^*-*^*/dcm*^*-*^ *E. coli* cells.

Even though a small increase in 5cadC was observed at higher enzyme concentrations (3 – 8 µM), the sum of 5hmdC and 5fdC remained stable at ∼95%, as the levels of 5mdC decreased with increasing enzyme concentrations.

We further investigated the effect of varying the concentration of the co-substrate α-ketoglutarate (αKG) on the catalytic activity of DM T940A/Y1567F Tet3. We tested αKG concentration ranging from 10 µM to 1000 µM. We noted that 1 mM αKG was optimal for achieving high combined levels of 5hmdC and 5fdC, which was once again accompanied by very low levels of 5mdC and 5cadC (Figure 2B).

Next, as oxidation reactions were routinely conducted for 1 h, the effect of extended incubation times on overall activity was evaluated by incubating gDNA with DM T940A/Y1567F Tet3 at 200 mM NaCl and 1 mM αKG for various time periods (Figure 2C). We observed that the sum of 5hmdC and 5fdC remained relatively stable after 1 h of reaction time, indicating that the maximum oxidation capacity is being gradually reached, since an increased treatment time did not result in considerably higher levels of 5cadC.

Lastly, we investigated the impact of the DNA amount supplied in the reaction on the distribution of oxidation products. To that end, 4 µM DM T940A/Y1567F Tet3 was reacted with genomic DNA ranging from 100– 1500 ng. We noted that all tested gDNA concentrations generated comparable amounts of 5hmdC and 5fdC. This suggests that under the applied reaction conditions (200 mM NaCl), further oxidation of fdC to cadC is largely restricted, irrespective of the amount of DNA to be oxidized. Taken together, we determined the best reaction condition for our purpose to be 4-8 µM DM T940A/Y1567F Tet3 applied to 1 µg genomic DNA in the presence of 200 mM NaCl and 1 mM αKG, incubated for 1 h.

Having now demonstrated that the DM T940A/Y1567F Tet3 stably produces >=95% 5hmdC and 5fdC over a wide range of enzyme concentrations and genomic DNA concentrations, at 200 mM NaCl, the question was raised as to whether different genomes containing varying levels of 5mdC would result in a similar distribution as observed for HEK293T gDNA. gDNA isolated from HEK293T cells typically contains ∼4.5% 5mdC per dG. Therefore, to account for any variability in the methylation levels, we oxidized genomic DNA isolated from mESCs, which were cultivated in the naïve (Figure 2E) and primed state (SI Figure 8), to assess the oxidation capacity at lower 5mdC levels compared to HEK293T gDNA. Approximately 0.81% and 2.99% of cytosines (per dG) are methylated in naïve and primed mESC gDNA, respectively. Conversely, to generate a gDNA substrate with substantially higher 5mdC levels, all CpGs in λ gDNA were methylated *in vitro* with the CpG-specific methyltransferase M.SssI, resulting in a genome in which 25.74% of all cytosines are methylated (Figure 2F). Quantitative MS analysis of the oxidation reaction revealed that the sum of 5hmdC and 5fdC is essentially comparable for all four types of gDNA substrates at enzyme concentrations of >= 4 µM. In all four gDNA substrates, 5mdC was oxidized to >=91% 5hmdC and 5fdC (HEK293T: >=95%, naïve mESCs: >=92%, primed mESCs: >=91%, Lambda DNA: >=95%), with minor amounts of 5mdC and 5cadC. These results demonstrate that conditions favoring the stable production of predominantly 5hmdC and 5fdC can be achieved and regulated with the ionic strength in the buffer and the amount of enzyme provided in the oxidation reaction. Furthermore, treatment of genomic DNA with DM-Tet3 did not yield an increase for both 8-oxo-2′-deoxyguanosine (8-oxo-dG) and 5-hydroxymethyl-2′-deoxyuridine (5hmdU) (SI Figure 9).

To validate that the ionic strength can regulate the distribution of the oxidation products, the catalytic efficiency of DM T940A/Y1567F Tet3 was determined using 5mdC-, 5hmdC-, and 5fdC-containing double-stranded oligonucleotide substrates. To that end, oxidation reactions were conducted at 100 mM and 200 mM NaCl at a constant enzyme concentration but with increasing amounts of the respective oligonucleotide (Figure 3). The conversion of 5mdC, 5hmdC, and 5fdC to the oxidation products 5hmdC, 5fdC, and 5cadC was analyzed by UHPLC-QQQ-MS/MS. The *in vitro* assays clearly demonstrate that DM T940A/Y1567F Tet3 exhibits higher enzymatic activity at 100 mM NaCl for all tested ds-DNA substrates (5mdC/5hmdC and f5dC) as previously observed in genomic DNA substrates (Figure 3). A distribution favoring the production of primarily 5hmdC and 5fdC was also observed at 200 mM NaCl. Thus, this confirms the selective and robust stalling of the oxidation at 5hmdC and 5fdC for the 5mdC containing ds-DNA oligonucleotide (Figure 3A). As expected, at 200 mM NaCl, the oxidation of 5fdC to 5cadC was strongly inhibited for the 5hmdC and 5fdC-containing ds-DNA oligonucleotide (Figure 3B and 3C). Overall, DM-Tet3 displayed very weak catalytic activity in oxidizing the 5fdC oligonucleotide at 200 mM NaCl (Figure 3C).

**Figure 3:**
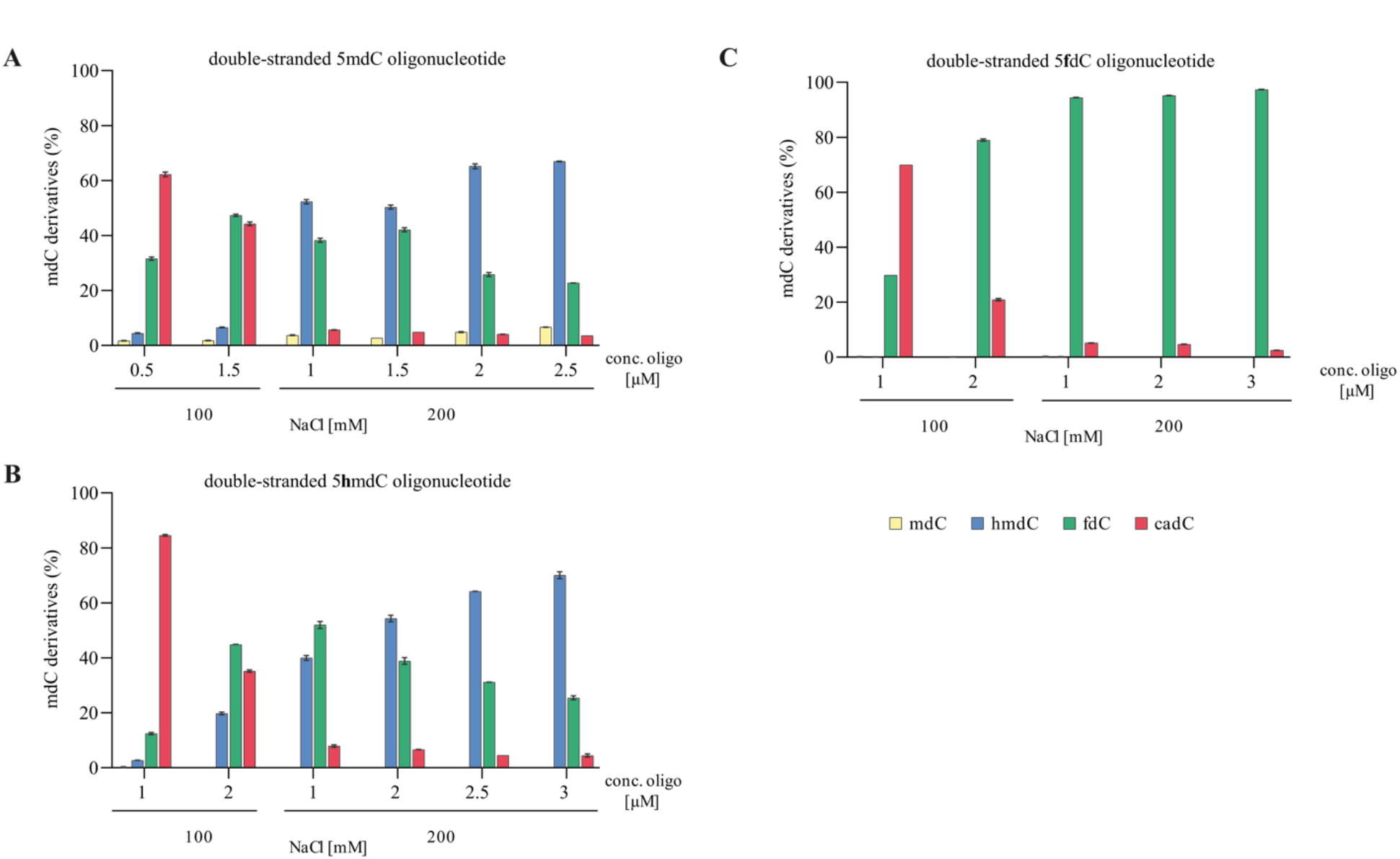
DM-Tet3 activity on non-canonical cytosines. Oxidation of (A) 5mdC, (B) 5hmdC, and (C) 5fdC in a double-stranded DNA oligo under distinct NaCl concentrations. All oxidation reactions were performed using 4 µM DM T940A/T1567F Tet3. Values represent mean ± SD, n =4.

Finally, in order to produce DNA that contains only 5hmdC, we aimed to chemically reduce the proportion of 5fdC, generated by our DM T940A/Y1567F Tet3, to 5hmdC. For this, we employed a previously reported protocol for the sequencing of 5fdC. In reduced bisulfite sequencing (redBS-Seq), Booth *et al*. reduce 5fdC using sodium borohydride (NaBH_4_).^19^

First, we performed M.SssI-catalyzed methylation of all CpGs within a DNA probe. To demonstrate the robustness and broad applicability, we prepared distinct DNA probes: A PCR amplicon, originating from a MeCP2 binding site in the Gabrb3 locus of the mouse genome (see SI Figure S10 and SI), a dam-/dcm-lambda DNA, and genomic DNA isolated from undifferentiated human iNGN cells. Subsequently, DNA was oxidized with DM T940A/Y1567F Tet3 and subjected to NaBH_4_ reduction (Figure 4A). The oxidation led to combined 5hmdC and 5fdC levels of 95.1 % for the lambda DNA (5hmdC: 63.4%, 5fdC: 31.7%), 92.9% for the MeCP2 binding site PCR product (5hmdC: 44.6%, 5fdC: 48.3%) and 93.4% for the genomic iNGN DNA (5hmdC: 44.0%, 5fdC: 49.4%) (Figure 4B). After reduction, 97.7% of 5hmdC for the lambda DNA, 93.7% for the PCR product, and 92.8% for the genomic iNGN DNA was achieved. Reversely, only low levels of 5mdC and 5cadC were observed (1-4% for each, respectively) (Figure 4B).

**Figure 4:**
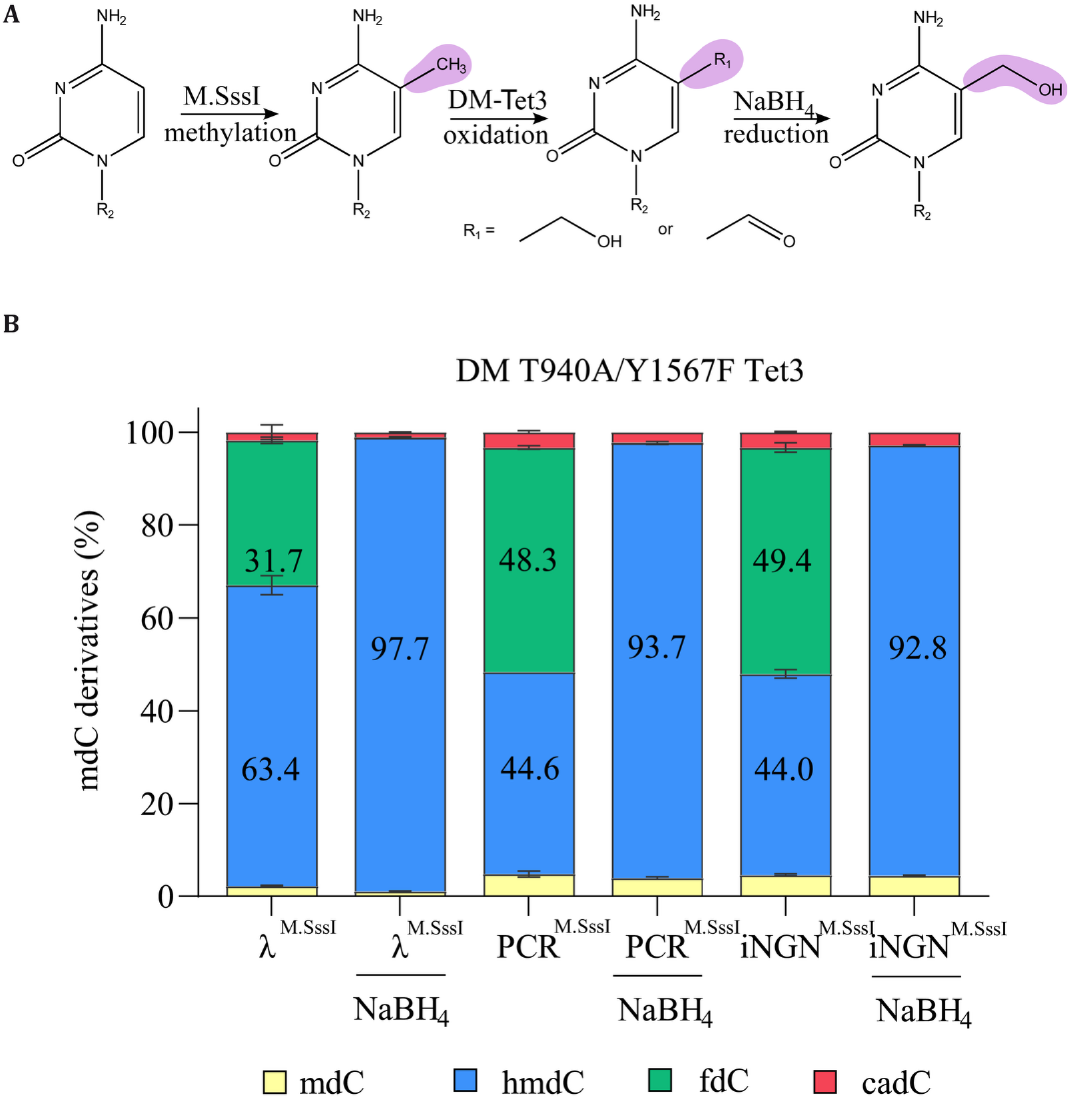
Generation of 5hmdC-DNA. (A) DNA methylation and oxidation scheme, DNA is first methylated using M.SssI, converting all cytosines in a CpG context to 5mdC; oxidized using the DM-Tet3 producing a mixture of 5hmdC and fdC; and eventually subjected to sodium borohydride reduction to convert all 5fdC to 5hmdC. (B) UHPLC-QQQ-MS/MS results of methylated-oxidized DNA as well as methylated-oxidized-reduced DNA – λ = Lambda DNA from *dam*^*-*^*/dcm*^*-*^ *E. coli* cells, PCR = a 531 bp PCR product, iNGN genomic DNA isolated from undifferentiated iNGN cells. Values represent mean ± SD, n =2.

## DISCUSSION

Modified nucleic acids are widely used in both research and medical diagnostics to study the role of modifications or as internal standards for diagnostic assays. In the case of 5hmdC, previously available approaches introducing 5hmdC were limited to generating large oligonucleotides (max. 100 bp–300 bp) or incorporating 5hmdC into native sequence CpG motifs.

We developed a Tet3 stalling mutant (DM-Tet3) that efficiently oxidizes 5mdC to 5hmdC and 5fdC, but only traces of 5cadC. Following subsequent sodium borohydride reduction of 5fdC, we generate DNA that almost exclusively contains 5hmdC. This approach allows the conversion of DNA of any size, including small synthetic oligos, PCR products, and even entire genomes, into 5hmdC. While this study is restricted to the use of M.SssI, other DNA methyltransferases can be used to generate 5hmdC in different sequence contexts, e.g., HpaII (CCGG). Moreover, through iterative use of methyltransferases, oxidation, and reduction, DNA with both 5mdC and 5hmdC can be created. This robust approach enables flexible and rapid production of 5hmdC-containing DNA with immediate implications for research and diagnostics.

Compared to previous techniques, the presented approach provides 5hmdC-containing DNA that accurately mimics the native, endogenous mammalian modification pattern. This allows precise investigation of how 5hmdC affects DNA-protein interactions or impacts chromatin formation. Additionally, due to the absence of length restrictions, the produced DNA resembles the biochemical characteristics of genomic DNA more closely. This is very valuable for NGS-based detection assays, which rely on chemical or enzymatic conversion of 5hmdC. Thus, these DNA fragments can serve as high-quality spike-ins or internal standards to improve the quantitative accuracy of NGS-based detection assays such as oxBS-seq or TAB-Seq. Lastly, direct sequencing technologies such as nanopore sequencing rely on pre-trained deep learning algorithms to infer DNA modifications on native DNA. While the models for detecting 5mdC are trained on M.SssI-based methylated DNA, the training of 5hmdC-detecting models is purely based on short synthetic oligos. The 5hmdC-containing DNA prepared here has the potential to improve the training of such models, leading to more sensitive and robust detection of 5hmdC across the genome.

Using 5hmdC as a biomarker in cancer and neurodegeneration requires high-quality standards. 5hmdC-containing DNA produced by our approach closely matches real biological DNA and thus represents ideal internal controls to validate diagnostic assays such as qPCR, digital PCR, and sequencing. This will improve the accuracy of the assays, prevent false negatives and false positives, and support inter-laboratory consistency.

In summary, our approach to generating 5hmdC-containing DNA using a Tet3 stalling mutant (DM-Tet3) and subsequent sodium borohydride reduction offers a robust and flexible method for producing DNA that closely mimics native mammalian modification patterns. This technique can potentially enhance biochemical assays for investigating the epigenetic role of 5hmdC in the mammalian genome. Furthermore, it may strengthen diagnostic assays such as qPCR, digital PCR, and NGS-based techniques by providing high-quality internal controls.

## Supporting information

SI1_Additional_Results

## RESOURCE AVAILABILITY

### Lead contact

Further information and requests for resources and reagents should be directed to and will be fulfilled by the lead contact, Pascal Giehr or Thomas Carell.

### Materials availability

All unique/stable reagents generated in this study are available from the lead contact with a completed materials transfer agreement.

### Data and code availability

Any additional information required to reanalyze the data reported in this paper is available from the lead contact upon request.

## ACKNOWLEDGMENTS

We thank CRC 1309 (ID: 325871075), CRC 1361 (ID: 393547839) and TRR 237 (ID: 369799452) for financial support. This project has received funding from the European Research Council (ERC) under the European Union’s Horizon 2020 research and innovation program under grant agreement No 741912 (EPiR) and under the Marie Skłodowska-Curie grant agreement No 861381 (Nature-ETN). Further support was obtained by the BMBF Cluster4Future program (Cluster for Nucleic Acid Therapeutics Munich, CNATM, ID: 03ZU1201AA) and by the intl. graduate program RNAmed (Elite network of Bavaria). Open Access funding enabled and organized by Projekt DEAL.

## AUTHOR CONTRIBUTIONS

Conceptualization, P.G., T.C., H.S.; methodology, H.S., S.I., A.L., I.P.; investigation, H.S., S.I., I.P.; writing - original draft, H.S., S.I., T.C. and P.G.; funding acquisition, T.C. and P.G.; resources, H.S. and S.I.; supervision, P.G. and T.C.

## DECLARATION OF GENERATIVE AI AND AI-ASSISTED TECHNOLOGIES

During the preparation of this work, the author(s) used Microsoft Copilot and Grammarly to correct spelling and grammar. After using this tool or service, the author(s) reviewed and edited the content as needed and take(s) full responsibility for the content of the publication.

