## Supplementary material for "Biochemical engineering of 5hmdC-DNA using a Tet3 double-mutant": SI1_Additional_Results

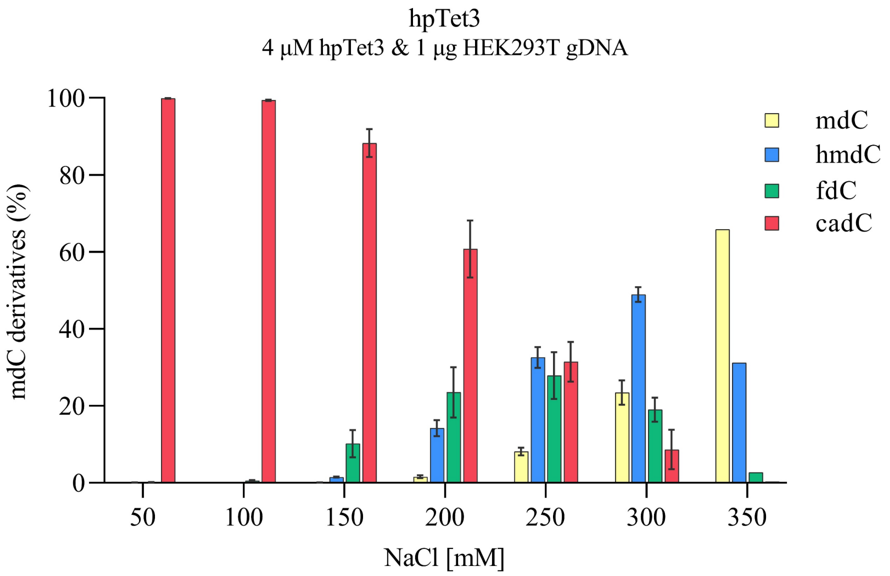


**Figure S1. Effect of different NaCl concentrations on the catalytic activity of hpTet3**. For each condition, 1 µg of HEK293T genomic DNA was incubated with 4 µM recombinant hpTet3 protein with the respective NaCl concentration at 37°C for 1 h. Samples were analyzed by UHPLC-QQQ-MS/MS. Values represent mean ± SD, n =3 for NaCl concentrations ranging from 50 mM–300 mM, n=1 for NaCl concentration of 350 mM.

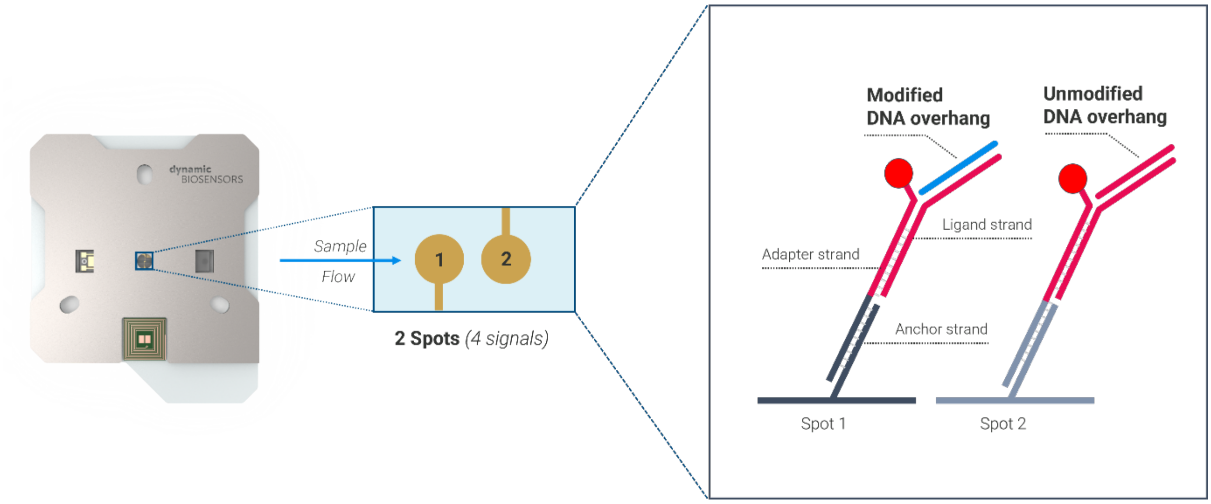


**Figure S2. heliX Adapter chip setup.** Binding kinetics are measured on two gold spots within the microfluidic channel of a heliX® chip. Graphical representation of Spot 1 used as measurement spot, in which the dsDNA overhang contains 5mdC modification, whereas Spot 2 is used as reference spot, with unmodified dsDNA overhang.

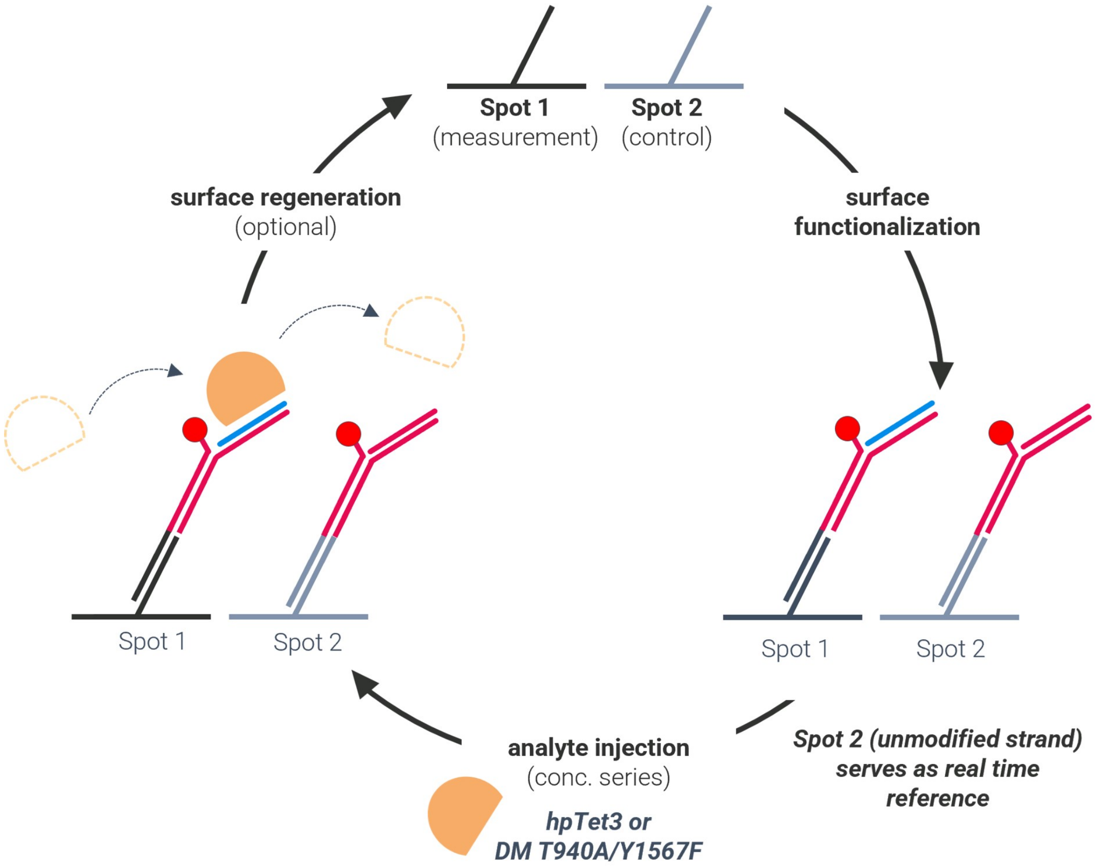


**Figure S3. Representation of the switchSENSE® experimental workflow**, consisting of 3 steps: (1) immobilization of the DNA ligands on the biochip surface; (2) Protein binding: association and dissociation kinetics are measured; (3) surface regeneration: injection of a high pH solution allows to remove the DNA ligands from the surface, and the biochip is ready to be functionalized again with fresh DNA ligand strands.

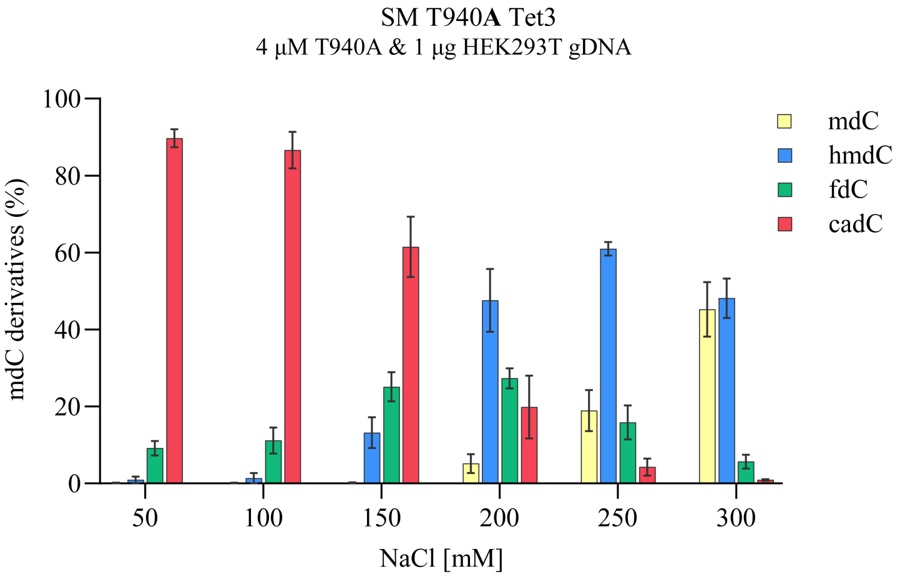


**Figure S4. Impact of different NaCl concentrations on the catalytic activity of SM T940A Tet3**. 1 µg of HEK293T genomic DNA was incubated with 4 µM recombinant SM T940A Tet3 protein with different NaCl concentrations at 37°C for 1 h. Samples were analyzed by UHPLC-QQQ-MS/MS. Values represent mean ± SD, n =3.

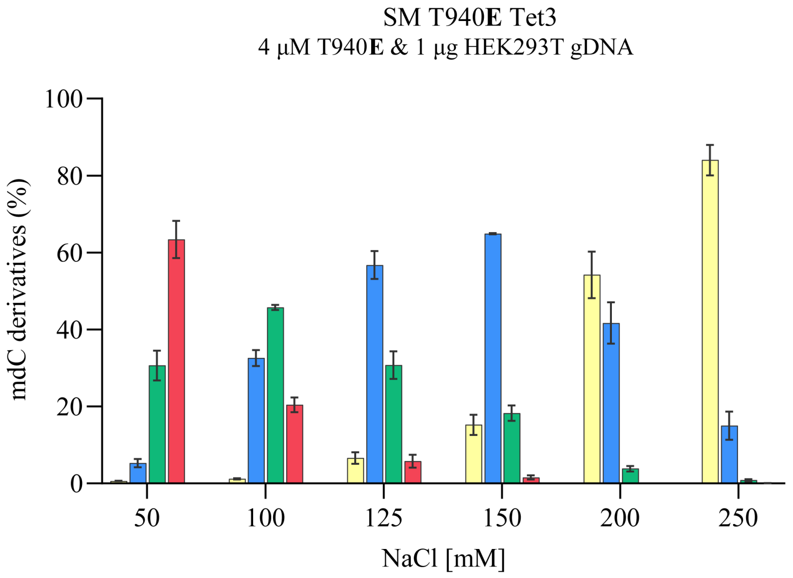


**Figure S5. Impact of different NaCl concentrations on the catalytic activity of SM T940E Tet3**. 1 µg of HEK293T genomic DNA was incubated with 4 µM recombinant SM T940E Tet3 protein with different NaCl concentrations at 37°C for 1 h. Samples were analyzed by UHPLC-QQQ-MS/MS. Values represent mean ± SD, n =4.

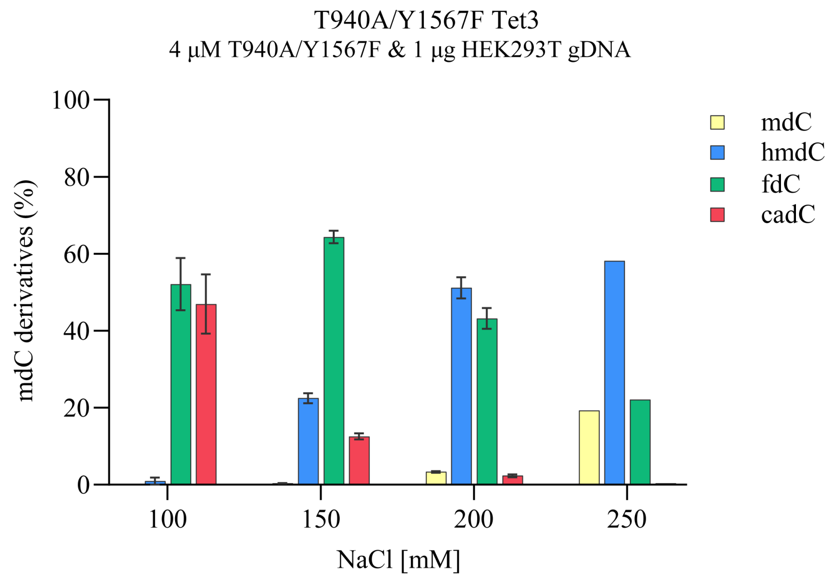


**Figure S6. Impact of different NaCl concentrations on the catalytic activity of DM T940A/Y1567F Tet3**. 1 µg of HEK293T genomic DNA was incubated with 4 µM recombinant DM T940A/Y1567F Tet3 protein with different NaCl concentration at 37°C for 1 h. Samples were analyzed by UHPLC-QQQ-MS/MS. Values represent mean ± SD, n =2 for NaCl concentrations of 100 mM, 150 mM, and 200 mM, n=1 for NaCl concentration of 250 mM.

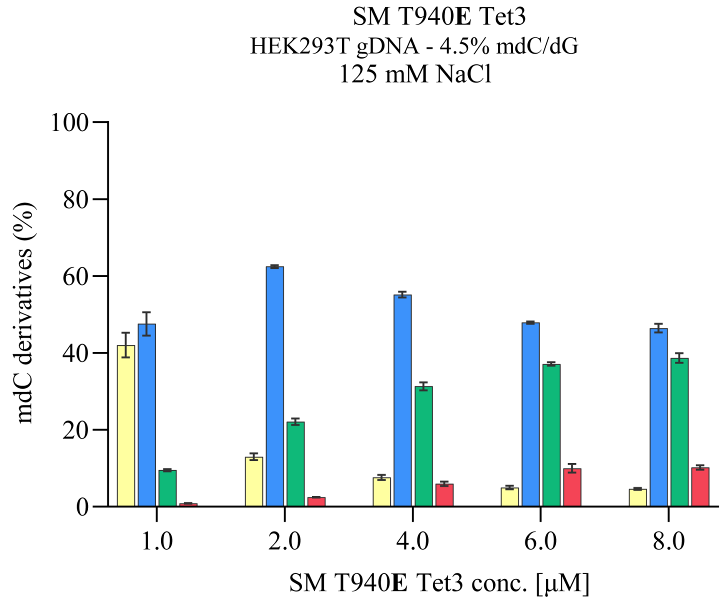


**Figure S7. Impact of varying SM T940E Tet3 concentrations on the oxidation of 5mdC to 5hmdC, 5fdC, and 5cadC at 125 mM NaCl.** For each reaction, 1 µg of human gDNA isolated from HEK293T cells was incubated with the respective amount of recombinant SM T1940E **Tet3** protein at 37°C for 1 h. Samples were analyzed by UHPLC-QQQ-MS/MS. Values represent mean ± SD, n =4.

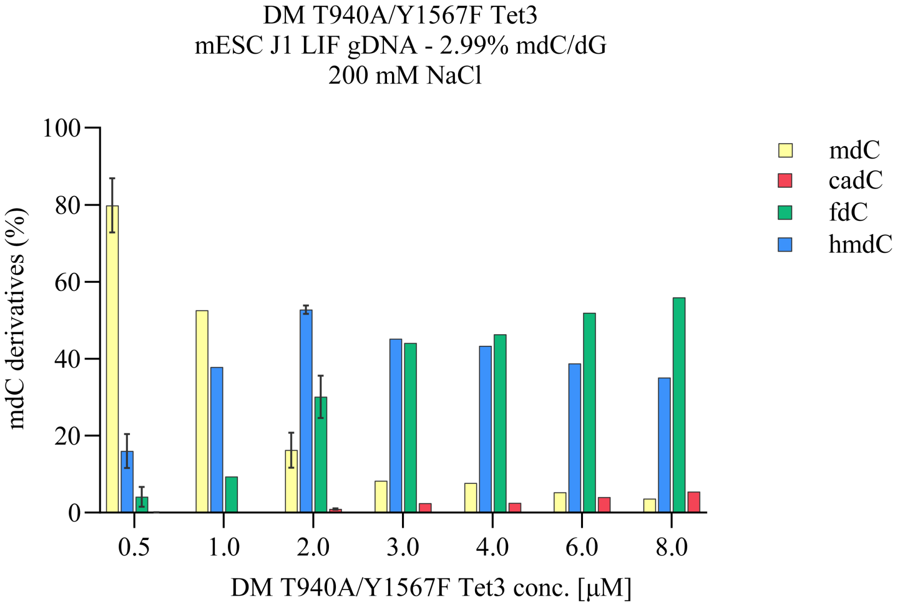


**Figure S8. Impact of various DM T940A/Y1567F Tet3 concentrations on the oxidation of 5mdC to 5hmdC, 5fdC, and 5cadC at 200 mM NaCl.** For each reaction, 1 μg of mouse gDNA isolated from J1 mouse embryonic stem cells cultivated in the primed state (mESC J1 LIF) was incubated with the respective amount of recombinant DM T940A/Y1567F Tet3 protein at 37°C for 1 h. Samples were analyzed by UHPLC-QQQ-MS/MS. Values represent mean ± SD, n =2 for 0.5 μM and 2 μM DM T940A/Y1567F Tet3, n=1 for the remaining DM T940A/Y1567F Tet3 concentrations.

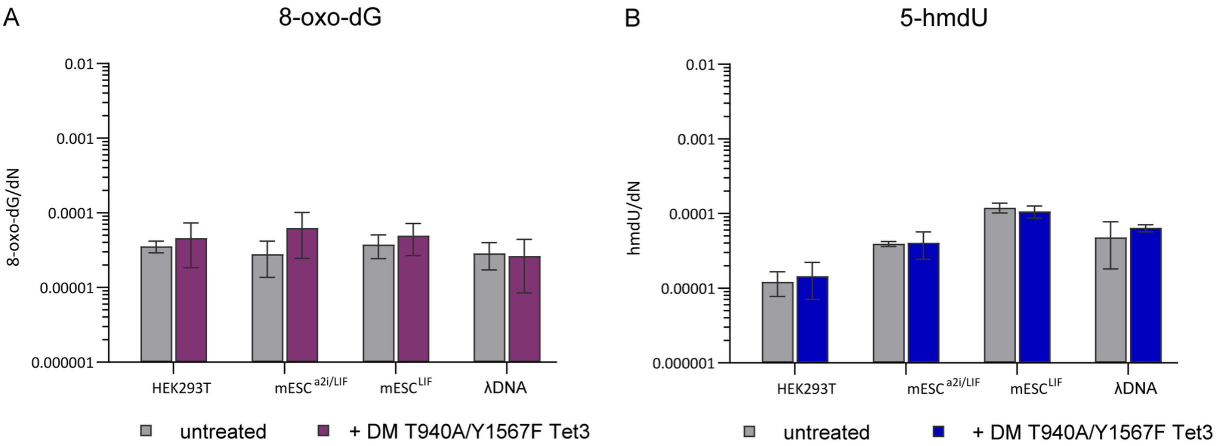


**Figure S9. 8-oxo-dG (A) and hmdU (B) levels per nucleosides (dN) of genomic DNA before and after treatment with DM T940A/Y1567F Tet3 as quantified by UHPLC-QQQ-MS.** Values represent mean ± SD, n =2.

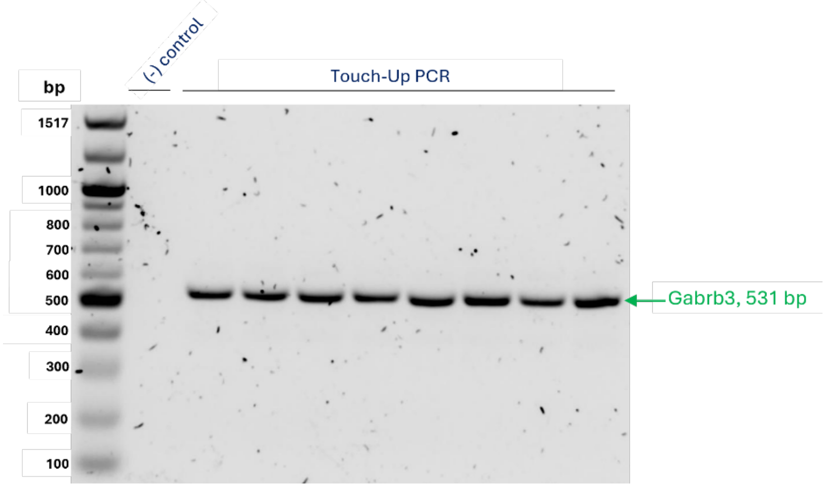


**Figure S10. Amplifications of the PCR product, Gabrb3, by Touch-Up PCR.** The PCR products were analysed on 1.2% agarose gels and compared with the 100 bp DNA Ladder (N3231L, New England Biolabs).

### Table S1. Kinetic Rates Overview.

| **hpTet3** | $k_{on1}$ (E+6 $M^{-1}s^{-1}$) | $k_{on2}$ (E+6 $M^{-1}s^{-1}$) | $k_{off1}$ (E-3 $s^{-1}$) | $k_{off2}$ (E-3 $s^{-1}$) | Rel. Amplitude $k_{off1}$, $k_{off2}$ (%) | $K_{d1}$ (nM) | $K_{d2}$ (nM) |
|---|---|---|---|---|---|---|---|
| HE140 | 4.02 ± 2.55 | 0.12 ± 0.01 | 121 ± 6 | 0.45 ± 0.03 | 50, 50 | 30.2 ± 19.2 | 3.89 ± 0.23 |
| HE200 | 0.32 ± 0.21 | **0.02 ± 0.01** | 159 ± 14 | **0.09 ± 0.04** | 35, **65** | 492 ± 323 | **4.00 ± 1.79** |
| HE300 | | | | N/D | | | |
| **DM** | | | | | | | |
| HE140 | 0.37 ± 0.09 | 0.37 ± 0.01 | 203 ± 13 | 0.84 ± 0.04 | 48, 52 | 552 ± 60.5 | 2.27 ± 0.12 |
| HE200 | 2.04 ± 1.57 | **0.08 ± 0.02** | 81 ± 16 | **0.74 ± 0.07** | 14, **85** | 39.8 ± 31.6 | **9.42 ± 2.38** |
| HE300 | | | | N/D | | | |
