## Supplementary material for "Biochemical engineering of 5hmdC-DNA using a Tet3 double-mutant": SI2_Methods

#### Expression and purification of *Mus musculus* hpTet3 and mutant variants in *E. coli*

Recombinant expression and purification of hpTet3 and the Tet3 mutant variants T940A, T940E, and T940A/1567F were performed as previously described for hpTet3 (Uniprot No. A0A5K1VVP6, aa 696 – 1604, with residues 1048 – 1508 replaced by a 15-residue GS-linker GGGGSGGGGSGGGGS).[1]

To incorporate the desired point mutation/s into the hpTet3 sequence for the mutant variants T940A, T940E, and T940A/Y1567F, DNA fragments with the corresponding point mutation were introduced by PCR and these mutation-containing fragments were cloned into the pET28a hpTet3 plasmid via Gibson assembly through overlapping primers using the NEBuilder® HiFi DNA Assembly Cloning Kit (New England Biolabs, Cat. No. E5520S). The identity of the sequences was confirmed by sequencing (Eurofins).

All proteins were expressed in *Escherichia coli* BL21 (New England Biolabs, Cat. No. C2527H). To that end, BL21 (DE3) competent E. coli cells were transformed with the corresponding plasmids and selected on LB plates containing kanamycin (25 µg/ml final concentration). A single colony was picked and cultured overnight (for 12–16 h) at 37 °C in 50 mL of LB broth (LB medium supplemented with 25 µg/ml kanamycin) at 180 rpm in an Innova S44i incubator 100 shaker (Eppendorf). The overnight culture was diluted 1:1000 in LB medium (supplemented with 25 µg/ml kanamycin) and incubated at 37 °C, 180 rpm until an $OD_{600}$ of ~0.3 was reached. Then, cultures were cooled down on wet ice to 16°C and protein expression was induced at an $OD_{600}$ between 0.5 and 0.6 by addition of isopropyl-β-D-thiogalactopyranoside (IPTG) to a final concentration of 0.5 mM. Additionally, cells were supplemented with a 1000x trace element stock solution to a concentration of 1x trace metals containing 50 µM $FeCl_2$, 10 µM $ZnSO_4$, 20 µM $CaCl_2$, 10 µM $MnCl_2$, 2 µM $CoCl_2$, 2 µM $CuCl_2$, 2 µM $NiCl_2$, 2 µM $Na_2MoO_4$, 2 µM $Na_2SeO_3$ and 2 µM $H_3BO_3$. IPTG-induced cultures were grown at 16 °C for 18 h. Cells were subsequently harvested by centrifugation (10 min at 8000 rpm at 4 °C), rinsed in cold PBS, and centrifuged once again at 8000 rpm. Cell pellets were flash-frozen in liquid nitrogen and kept at −80 °C until purification. The frozen cell pellets were thawed on ice, resuspended in ice-cold lysis buffer (50 mM HEPES, 500 mM NaCl, 10% v/v glycerol, 10 µM $ZnCl_2$ mM, 5 mM $MgCl_2$, 5 mM ATP, 0.5 mg/mL lysozyme, 0.5 mM tris(2-chloroehyl)phosphate (TCEP), pH 6.8) supplemented with Benzonase (Merck, Cat. No. 1016540001) and cOmplete™, EDTA-free Protease Inhibitor Cocktail (Roche, Cat. No. 11873580001) and incubated on ice for 30 min. The cell suspension was lysed by homogenization using a high-pressure homogenizer (EmulsiFlex C5, Avestin Inc.) at 400 mbar for three rounds. Cleared lysate was obtained by centrifugation at 40,000 g for 45 min at 4 °C. The supernatant was transferred to a new tube, and centrifugation was repeated for another 15 min (40,000 g, 4°C). The cleared lysate was filtered and loaded onto a 5 ml StrepTrap XT column (Cytiva, Cat. No. 29401322) using a 1 ml/min flow rate. After sample application, the column was washed with 8 column volumes (CV) of wash buffer (50 mM HEPES, 500 mM NaCl, 10% v/v glycerol, 0.5 mM TCEP, pH 6.8) until a stable baseline was reached and bound protein was eluted with 10 CV elution buffer (50 mM HEPES, 100 mM NaCl, 0.5 mM TCEP, 10% v/v glycerol, 50 mM biotin, pH 6.8) using a flow-rate of 3 mL/min. Protein-containing fractions were pooled and concentrated using Amicon Ultra 15 mL centrifuge filter units (30 kDa molecular cutoff). The concentrated eluate was diluted 1-fold in ATP buffer (50 mM HEPES, 100 mM NaCl, 0.5 mM TCEP, 10% v/v glycerol, 10 mM $MgCl_2$, 10 mM ATP, pH 6.8) to remove tightly bound molecular chaperone Hsp70 (DnaK) – Hsp40 (DnaJ) – hpTet3 or mutant protein complexes and incubated for 30 min on ice. Incorrectly folded protein aggregates were removed by centrifugation for 30 min at 25,000 g at 4°C. Then, the supernatant was loaded onto a 5 ml HiTrap Heparin HP column (Cytiva, Cat. No. 17040701) pre-equilibrated in binding buffer (50 mM HEPES pH 6.8, 0.1 M NaCl, 0.5 mM TCEP, 10% v/v glycerol) and the column was washed with 20 CV of binding buffer. The protein was eluted with a linear salt gradient over 20 CV (0%–100% elution buffer (50 mM HEPES pH 6.8, 1.5 M NaCl, 0.5 mM TCEP, 10% v/v glycerol) starting from 0.1 to 1.5 M NaCl). hpTet3 and all mutant variants elute around 300 mM NaCl.

Pure protein-containing fractions were pooled, buffer exchanged to a final buffer containing 50 mM HEPES pH 6.8, 300 mM NaCl, 10% glycerol, and 0.5 mM TCEP, and concentrated using Amicon centrifugal filters with an MWCO of 30 kDa. Aliquots of pure protein were flash-frozen in liquid nitrogen and stored at -80 °C. Typical yields were 2-3 mg per liter of *E. coli* culture.

All protein purification steps were carried out at 4 °C with pre-cooled buffers on a ÄKTA pure chromatography system (Cytiva) and protein-containing fractions were kept on ice during whole purification procedure.

**Preparation of the PCR product**

We designed an amplicon across a Mecp2 binding region based on the ChIP-Seq data from Baubec *et al.*[2]

***Table S2:*** *Gabrb3-Amplicon Location and Sequence.*

| PCR Product | Sequence 5'-3' (UCSC, mm9) |
|---|---|
| Gabrb3, 531 bp<br>mm9, chr7: 64,846,301-64,846,837 | TGTCCAGCGCGGATCTGCGGCCGCACGCAAGCCCTTCCCC<br>GGGAGTCCGGTGCGGCCGCTGCCGCGGCTTGGGAGGGCG<br>CCGGCGCCCCGGAGCGACCGCGGACCCATAGGGGGCGGG<br>TGCTGTTCTTGTCCGGCTCCGCCCCCGCCTCCGCCCGCCCA<br>GCGCACCCGCGCCAGGGCTCCTCCCTCCCGGCCCGCTCCT<br>CCCCCCTGCTCTCCTCCCCCTCCCCCTCCGCCTCCTCCGCT<br>CCGGGCCAGCGCGGCGGCGGTGGCGGCAGCAGCAGCAGC<br>AGCCCCGGCTGCGGGTCGCGACGGCGGCGGGGCGCCCCC<br>TCCCCCGTGCCGGGGCGCGGCGAAGGGATGTGGGGCTTTG<br>CGGGAGGAAGGCTTTTCGGCATCTTCTCGGCCCCGGTGCTG<br>GTGGCGGTGGTTTGCTGCGCTCAGAGGTAGGGTCGCTGGT<br>GGGTTGACGGGAAGCGGGCCGGGCGCGAGCTGGCGCGCT<br>GTGTGCGCGCCGGGCGCTGGCGGGTGAGCCGCTGCCTGAC<br>CCTGTTCTCTGTGCCATGT |

Primers were designed manually using the free SnapGene Viewer version **8.0.3** [3] and were purchased from **Microsynth AG**.

***Table S3:*** *Primers for PCR reactions.*

| ***Primer name*** | ***Primer Sequence 5'-3'*** | ***Primer size*** |
|---|---|---|
| **Gabrb3-FWD** | 5'-TGT CCA GCG CGG ATC TGC GG-3' | 20nt |
| **Gabrb3-REV** | 5'-CAC AGA GAA CAG GGT CAG GCA GCG-3' | 24nt |

PCR was conducted using Q5® Hot Start High-Fidelity 2X Master Mix (M0494S, NEB):

***Table S4:*** *PCR reaction components.*

| Component | 25 µl Reaction | Final Concentration |
|---|---|---|
| Q5 High-Fidelity 2X Master Mix | 12.5 µl | 1X |
| 10 µM Forward Primer (Gabrb3) | 1.25 µl | 0.5 µM |
| 10 µM Reverse Primer (Gabrb3) | 1.25 µl | 0.5 µM |
| Genomic Mouse DNA, 4.05 ng/µl | 4.94 µl | 20 ng (< 1,000 ng) |
| Nuclease-Free Water | 5.06 µl | |

To establish optimal annealing conditions, the standard Q5 Hot Start High-Fidelity 2X Master Mix PCR protocol was modified using a Touch-Up PCR strategy. The melting temperature (Tm) of the primers was calculated using the NEB Tm Calculator version 1.16.7 [4], yielding a value of 72 °C. Thermal cycling was initiated at an annealing temperature of 68°C, approximately 4°C below the calculated Tm, and incrementally increased by 1°C per cycle over the first five cycles. This gradual increase was followed by 30 amplification cycles conducted under the standard conditions recommended by the manufacturer.

***Table S5****: Thermocycler program of the Gabrb3 Touch-Up PCR*

| STEP | TEMP | TIME |
|---|---|---|
| Initial Denaturation | 98°C | 30 seconds |
| 1–5 Cycles | 98°C | 10 seconds |
| Denaturation, annealing & elongation | 68°C, with a 1°C increase after each cycle (68–72°C) | 30 seconds |
| | 72°C | 20 seconds |
| 5–35 Cycles | 98°C | 10 seconds |
| Denaturation, annealing, and | 72°C | 30 seconds |
| elongation | 72°C | 20 seconds |
| Final Extension | 72°C | 2 minutes |
| Hold | 4–10°C | ∞ |

Gabrb3 PCR products were purified using 1.2x MagMax™ Pure Bind Beads (ThermoFisher, Cat. No. A58522). Beads were added directly to the PCR reaction, incubated for 5 min at room temperature, and washed with freshly prepared 80% ethanol. Washing was repeated once for a total of two wash steps. Ethanol was removed, and the beads were dried for 1 minute with the lid open. The Gabrb3 PCR product was eluted with prewarmed (50°C) double-distilled water for 5 min in a thermocycler at 37°C. Figure S10 (SI additional results) depicts the results of the amplification of the PCR product, Gabrb3, by Touch-Up PCR.

**Preparation of methylated Gabrb3-Amplicon**

CpG methylation of the PCR product was performed in vitro using 400 ng of PCR product and 4 U of M.SssI enzyme (NEB, cat. no. M0226S) in the presence of 160 µM SAM in the provided reaction buffer (10 mM Tris-HCl pH 7.9, 50 mM NaCl, 10 mM MgCl2 and 1 mM DTT) in a total volume of 50 µL for 120 min at 37°C. After 120 min, for complete CpG methylation, the reaction mixture was further supplemented with additional reaction buffer (0.1x), 145 µM SAM, and 4U M.SssI in a 55 µL final reaction volume and incubated for an additional 120 min at 37°C, followed by a heat inactivation for 20 min at 65°C. Methylated PCR products were purified using 1.2x MagMAX™ Pure Bind Beads (ThermoFisher, Cat. No. A58522) as described above. Quantitative UHPLC-QQQ-MS/MS confirmed the efficiency of DNA methylation.

**Preparation of methylated DNA**

CpG methylation of the different DNA substrates (lambda DNA (dam−, dcm−) (Thermo Scientific, cat. no. SD0021) and iNGN DNA) was performed in vitro using 1 µg of lambda DNA and iNGN DNA, and 4 U of M.SssI enzyme (NEB, cat. no. M0226S) in the presence of 160 µM SAM in the provided reaction buffer (10 mM Tris-HCl pH 7.9, 50 mM NaCl, 10 mM $MgCl_2$ and 1 mM DTT) in a total volume of 50 µL for 120 min at 37°C. After 120 min, for complete CpG methylation, the reaction mixture was further supplemented with additional reaction buffer (0.1x), 145 µM SAM, and 4U M.SssI in a 55 µL final reaction volume and incubated for an additional 120 min at 37°C, followed by a heat inactivation for 20 min at 65°C. Methylated lambda DNA and iNGN DNA were purified using 1.8x AMPure XP beads (Beckman Coulter, product no. A63881). Beads were directly added after M.SssI inactivation; the reaction was mixed by pipetting followed by an incubation for 20 min at room temperature on a Hulamixer. Liquid was briefly centrifuged and incubated on a magnetic stand until the liquid became clear. After discarding the supernatant, the beads were washed twice with 500 µl freshly prepared 80% ethanol. Beads were dried for 1 minute at room temperature while still on the magnetic stand, and the DNA was eluted with prewarmed (50°C) nuclease-free water for 30 min at 37°C. The efficiency of DNA methylation was confirmed by quantitative UHPLC-QQQ-MS/MS.

**Oxidation reactions with hpTet3 and the different mutant variants for salt dependency experiments**

Oxidation reactions were performed at a final volume of 50 µl at 37 °C for 1 h at 550 rpm in a thermomixer. DNA concentrations were determined using the Qubit dsDNA 1x HS-Assay-Kit (ThermoFisher, cat. no. Q32851).

For salt dependency experiments in Figure 1B, all reactions were performed with 1 µg of genomic DNA isolated from HEK293T cells. To that end, a 2X master mix was prepared, which contained 100 mM HEPES pH 7.9, 2 mM α-ketoglutarate, 4 mM ascorbic acid, 2.4 mM ATP, and 5 mM DTT. Additionally, to this 2x master mix, sodium chloride (NaCl) to a final concentration of 140 mM or 340 mM NaCl was added. In this manner, NaCl concentrations of 70 mM or 170 mM NaCl in the oxidation reaction volume of 50 µl were obtained, respectively. Since all enzymes were stored in a storage buffer which contained 300 mM NaCl (storage buffer composition: 50 mM HEPES pH 6.8, 300 mM NaCl, 10% glycerol, 0.5 mM TCEP), 1/10 (5µl) of the final reaction volume (50 µl) was used to add the respective enzymes diluted in storage buffer to a final concentration of 4 µM (10 µg). Adding 1/10 of the final reaction volume (50 µl) accounted for an additional 30 mM NaCl in the total reaction and yielded final NaCl concentrations of 100 and 200 mM NaCl. This aimed to have the same NaCl concentrations for all tested enzymes by controlling and compensating for fluctuations in the enzyme concentration. The pH of the reaction mix was 7.4 after the addition of all buffer components. The reaction was initiated by adding 105 µM $Fe(NH_4)_2(SO_4)_2$ and incubated at 37 °C for 1 h. Subsequently, 4 U of Proteinase K and SDS to a final concentration of 0.2 % were added to the reaction mixture and incubated for 1h at 50 °C. Oxidized gDNA was purified with 1.8x AMPure XP beads (Beckman Coulter, product no. A63881) according to the optimized protocol described above. Oxidation efficiency was quantified by UHPLC-QQQ-MS/MS. Ascorbic and $Fe(NH_4)_2(SO_4)_2$ were freshly prepared and Fe(II) was dissolved in water and added immediately before the reaction begins to minimize the oxidation to Fe(III).

All oxidation reactions at different NaCl concentrations ranging from 50 mM to 350 mM NaCl (see additional results Figures S1, S4-6) were performed as described above using the 2x master mix with the corresponding NaCl concentrations.

**Oxidation reactions using double-mutant T940A/Y1567F Tet3 at 200 mM NaCl**

For reactions at 200 mM NaCl, a 2X master mix was prepared which contained 100 mM HEPES pH 7.9, 340 mM NaCl, 2 mM α-ketoglutarate, 4 mM ascorbic acid, 2.4 mM ATP and 5 mM DTT.

The DNA was added to this 2x master mix and analogous to the salt dependency experiments, for all oxidation reactions at 200 mM NaCl using the double-mutant (DM) T940A/Y1567F Tet3 enzyme, 1/10 (5µl) of the final reaction volume (50 µl) was used to add the corresponding amount of DM T940A/Y1567F Tet3 enzyme diluted in storage buffer to always have the same NaCl concentration of 200 mM NaCl in all reactions. This once again accounted for an additional 30 mM NaCl. After adding the enzyme/storage buffer mixture, the oxidation was initiated with the addition of 105 µM $Fe(NH_4)_2(SO_4)_2$ and incubated at 37 °C for 1 h at 550 rpm. After that, 4 U of Proteinase K (NEB, cat. no. P8107S) and SDS to a final concentration of 0.2% were added to the reaction mixture and incubated for 1h at 50 °C. Subsequently, oxidized lambda DNA and iNGN DNA were purified with 1.8x AMPure XP beads (Beckman Coulter, product no. A63881) according to the optimized protocol described above. Oxidized PCR products were purified using 1.2x MagMAX™ Pure Bind Beads (ThermoFisher, Cat. No. A58522) according to the optimized protocol described above.

For the DM T940A/Y1567F Tet3 titration experiment in Figure 2A, once again, 1/10 (5 µl) of the final reaction volume (50 µl) was used to add the respective DM T940A/Y1567F Tet3 enzyme amount diluted in storage buffer (ranging from 0.5 µM to 8 µM).

αKG (Figure 2B) and genomic DNA titration (Figure 2D) reactions were performed with the respective αKG and gDNA amounts as described above. For time point reactions in Figure 2, the reaction was further stopped by adding 5 mM EDTA.

The following DM T940A/Y1567F Tet3 final concentrations, depending on the methylation level for an optimal formation of >=92% 5hmdC and 5fdC, were used for the oxidation of 1 µg gDNA: **HEK293T** (4.5% mdC/dG): 6 µM (>=95% 5hmdC and 5fdC); **iNGN M.SssI** (6.31% mdC/dG): 8 µM (>=93% 5hmdC and 5fdC); **Lambda DNA M.SssI**: (25.74% mdC/dG): 8 µM (>=95% 5hmdC and 5fdC); **PCR product M.SssI** (33.82% mdC/dG): 8 µM (>=93% 5hmdC and 5fdC)

In detail oxidation reactions were performed as follows:

1. Prepare a 2x TET buffer

   2x TET buffer
   100 mM HEPES, pH 7.9
   340 mM NaCl
   2 mM α-ketoglutarate
   4 mM ascorbic acid (freshly prepared)
   2.4 mM ATP
   5 mM DTT

2. Prepare a fresh 2.625 mM $Fe(NH_4)_2(SO_4)_2$ stock solution (25x)

3. After dissolving $Fe(NH_4)_2(SO_4)_2$ in water, quickly assemble the reaction in the following order:

| | |
|---|---|
| 25 µl | 2x TET buffer |
| X.X µl | $H_2O$ (up to 50 µl) |
| 1 µg | genomic DNA |
| 5 µl | 10 – 22 µg/4 – 8 µM DM T940A/Y1567F Tet3 diluted in storage buffer |
| 2 µl | 25x $Fe(NH_4)_2(SO_4)_2$ |

   Incubate at 37°C for 1 hour at 550 rpm.

**Sodium borohydride reduction of double-mutant T940A/Y1567F Tet3 oxidized samples**

Sodium borohydride reduction reactions of double-mutant (DM) T940A/Y1567F Tet3 oxidized samples were performed in a final volume of 40 µl at room temperature in the dark. To that end, fresh sodium borohydride (Sigma, cat. no. 213462) solution (1 M) in water was prepared, and 10 µl of this freshly prepared 1 M $NaBH_4$ solution was added to 30 µl of 1 µg of DM T940A/Y1567F Tet3 oxidized DNA. The reaction mixture was vortexed and centrifuged, then held at room temperature (25°C) in the dark for 1 h. The lids were kept open for the whole reaction time to release pressure from gas generation, and every 15 min, reactions were vortexed and centrifuged to remove bubbles formed due to gas generation. Then, the reaction was quenched slowly by adding 20 µL of 750 mM sodium acetate (Sigma, cat. no. S2889) solution (pH 5). Upon addition, a violent release of hydrogen gas occurred. The reaction was held at room temperature for 50 minutes (≥10 minutes or until no further gas was released), in between the reaction was vortexed and centrifuged after a short interval (after 10-15 minutes each time), to remove the bubbles generated from gas generation. Reduced gDNA was purified with 1.8x AMPure XP beads (Beckman Coulter, product no. A63881) according to the optimized protocol described above. The reduction samples from the PCR products were purified using 1.2x MagMAX™ Pure Bind Beads (ThermoFisher, Cat. No. A58522) according to the optimized protocol described above.

**Synthesis of modified oligonucleotides**

5-hydroxymethyl-2'-deoxycytidine and 5-formyl-2'-deoxycytidine containing oligonucleotides used in Figure 3 were synthesized according to the literature. [5]

**Cell culture**

HEK293T, iNGN, and mESCs were used in this study. The cell lines were cultivated at 37 °C in water-saturated, $CO_2$-enriched (5%) atmosphere.

HEK293T cells (CLS) were maintained in Dulbecco's Modified Eagle's Medium (DMEM) with high glucose content (Sigma-Aldrich D6546), supplemented with 10% (v/v) fetal bovine serum (FBS) (Life Technologies 10500-064), 1% (v/v) L-alanyl-L-glutamine (Sigma-Aldrich G8541), and 1% (v/v) penicillin–streptomycin (Sigma-Aldrich P0781). The cells were routinely passaged at a ratio of 1:10 when a confluence of 70–80% was reached.

J1 mESCs were cultured and maintained as previously reported in naïve state on 0.2% (w/v) gelatine-coated plates in DMEM (Sigma-Aldrich D6546), supplemented with 10% (v/v) Pansera ES-grade FBS (Pan Biotech), 1× MEM-nonessential amino acids (NEAA, Sigma-Aldrich M71145), 2 mM L-alanyl-L-glutamine, 1x Penicillin-Streptomycin (Sigma-Aldrich AP078), 0.1 mM β-mercaptoethanol, 103 U/mL mouse recombinant leukemia inhibitory factor (mLIF, Sigma-Aldrich ESG1107), 1.5 µM CGP 77675 (Sigma-Aldrich SML0314) and 3 µM CHIR 99021 (Axon Medchem) (a2iL conditions). In the naïve state, cells were passaged every 2 – 3 days in a ratio of 1:4 to 1:8 when a confluency of 60 – 75% was reached. For the naïve-to-primed mESC transition, cells were grown in medium supplemented with the components described above but without the addition of the GSK3α/β and Src kinase inhibitors. Cells were primed for 72 h in total before gDNA isolation.

iNGNs were grown on Geltrex (ThermoScientific, Cat. No. A1413201)-coated tissue plates. Coating solution was prepared by thawing Geltrex on ice and diluting 1:1000 in ice-cold DMEM/Ham's F-12 (F-12, Sigma-Aldrich, Cat. No. N4888) to a final concentration of 15 µg/mL, and plates were incubated for at least 2 h in the incubator at 37°C. iNGNs were cultured in the undifferentiated state in a medium containing 1:1 DMEM: F-12, 2 mM L-alanyl-glutamine, 0.1 mg/mL penicillin-streptomycin, 0.2 mM L-ascorbic acid 2-phosphate, 77.6 nM sodium selenite, 10.90 mM NaCl, 10 µg/mL hHOLO-Transferrin (Merck Millipore, Cat. No. 616424), 20 µg/mL hrInsulin (Sigma Aldrich, Cat. No. I9278), 100 ng/mL hrFGF-2 (PeproTech AF-100-18B), 2.0 ng/mL hrTGF-β1 (PeproTech, Cat. No. 100-21C). When a confluency of 60% was reached, iNGNs were passaged at a ratio of 1:4. Additionally, when passaging, 2 µM (final concentration) of ROCK inhibitor - Thiazovivin (Merck Millipore, Cat., No. 420220) was added to the medium.

**Isolation of gDNA**

gDNA was isolated using a previously published method by Traube *et al.* [6] Cells were washed with PBS and lysed directly in the plates with RLT buffer (Qiagen) supplemented with 0.01 equivalents of 2-Mercaptoethanol (14.3 mM final concentration), antioxidants 3,5-di-tert-butyl-4- hydroxytoluene (BHT, 200 μM) and deferoxamine mesylate salt (Desferal, 200 μM). To homogenize the lysate and shear the gDNA, samples were further subjected to bead milling using a Qiagen TissueLyser for 30s at 30 Hz. After cell lysis, gDNA isolation was performed as described in Traube *et al.* [6] Isolated gDNA was subjected to nucleoside digestion and UHPLC-QQQ-MS/MS measurement before and after oxidation.

**DNA digestion for LC–MS/MS**

gDNA samples or oligonucleotides before and after oxidation with the respective proteins were digested to nucleosides using the Nucleoside Digestion Mix from NEB (M0649S) in a total volume of 50 μL using 1 μL of enzyme and 5 μL of 10x reaction buffer at 37°C for 3 h. After 3 h, digested samples were filtered using an AcroPrep Advance 96-well Supor filter plate, 0.2 μm (Pall Life Sciences, (Pall, cat. no. 518-0022) and subjected to UHPLC-QQQ-MS/MS.

**UHPLC-QQQ-MS of nucleosides**

Canonical and modified nucleosides were quantified using the previously published stable isotope dilution LC-MS technique by Traube *et al.*[6] For the absolute quantification of nucleosides, an Agilent 1290 Infinity II equipped with a variable wavelength detector (VWD) combined with an Agilent Technologies G6490 Triple Quad LC/MS system with electrospray ionization (ESI-MS, Agilent Jetstream) was used. Nucleosides were separated using a Poroshell 120 SB-C18 column (2.7 μm, 2.1 × 150 mm; Agilent Technologies, cat. no. 683775-902) or a Poroshell 120 SB C8 column (2.7 μm, 2.1 × 150 mm; Agilent Technologies, cat. no. 683775-906) at 35 °C and a flowrate of 0.35 mL/ min using water supplemented with 0.0075% (vol/vol) formic acid (FA) (buffer A) and acetonitrile (MeCN) supplemented with 0.0075% (vol/vol) FA (buffer B). The gradient started with 100% buffer A, followed by an increase to 3.5% buffer B over a period of 4 min. Buffer B was then increased to 5% over 3 min from 4 min to 7 min. Then, from 7.0 min to 7.5 min, buffer B was increased further to 80 % and maintained at 80 % for 2.0 min before returning to 100 % solvent A in 0.5 min, and lastly a 3.0 min re-equilibration period. For the ionization of the nucleosides an ESI source was used. The N2 gas temperature was 120 °C with a flow rate of 11 L/min. Sheath gas temperature was 280 °C with a flow rate of 11 L/min. Capillary voltage was 3000 V, nozzle voltage was 0 V, nebulizer pressure was 60 psi, high-pressure RF at 150 V, and low-pressure RF at 60 V. The cell accelerator voltage was 5 V. The instrument was operated in dynamic MRM and positive-ion mode. The fragmentor voltage was 380V for all compounds. Other compound-dependent parameters are summarized in Table S6. For absolute quantification, each sample was co-injected with 1 μL of 0.5 μM stable isotope-labeled internal standard (ISTD) mix containing the following isotope standards: [$^{15}N_5$-$^{13}C_{10}$]-dA, [$^{13}C_9$]-dC, [$^{15}N_5$-$^{13}C_{10}$]-dG, [$^{15}N_2$-$^{13}C_{10}$]-dT, [$D_3$]-5mdC, [$D_2$-$^{15}$N2]-5hmdC, [$^{15}N_2$] 5fdC, [$^{15}N_2$]-5cadC, [$^{15}N_5$]-8-oxo-dG, and [$D_2$]-hmdU. For calibration, solutions of synthetic nucleoside standards were used, and analogous to the samples, 1 μL of 0.5 μM stable ISTD mix was co-injected with each calibration. Calibration curves and the sample data were analyzed using Agilent's Quantitative MassHunter Software (v B07.01) using the built-in calibration function.

**Table S6**: Compound-dependent LC-MS/MS parameters. $R_t$: retention time, CE: collision energy, CAV: collision cell accelerator voltage. $R_t$ values are given for the Poroshell 120 SB C8 column.

| Compound | Precursor ion (m/z) | MS1 resolution | Product ion (m/z) | MS2 resolution | $R_t$ (min) | CE (V) | CAV (V) | Polarity |
|---|---|---|---|---|---|---|---|---|
| **[$^{15}N_2$]-5cadC** | 274.08 | Wide | 158.03 | Unit | 2.7 | 6 | 5 | Positive |
| **5cadC** | 272.09 | Wide | 156.04 | Unit | 2.7 | 6 | 5 | Positive |
| **[$D_2$-$^{15}N_2$]-5hmdC** | 262.12 | Wide | 146.07 | Unit | 1.9 | 4 | 5 | Positive |
| **5hmdC** | 258.11 | Wide | 142.06 | Unit | 1.9 | 4 | 5 | Positive |
| **[$D_3$]-5mdC** | 245.13 | Wide | 129.09 | Unit | 2.5 | 4 | 5 | Positive |
| **5mdC** | 242.11 | Wide | 126.07 | Unit | 2.5 | 4 | 5 | Positive |
| **[$D_2$]-5hmdU** | 261.08 | Wide | 145.1 | Unit | 3.5 | 4 | 5 | Positive |
| **5hmdU** | 259.08 | Wide | 143.1 | Unit | 3.5 | 4 | 5 | Positive |
| **[$^{15}N_5$]-8-oxo-dG** | 289.09 | Wide | 173.04 | Unit | 6.0 | 9 | 5 | Positive |
| **8-oxo-dG** | 284.1 | Wide | 168.05 | Unit | 6.0 | 9 | 5 | Positive |
| **[$^{15}N_2$]-5fdC** | 258.09 | Wide | 142.04 | Unit | 5.2 | 5 | 5 | Positive |
| **5fdC** | 256.09 | Wide | 140.05 | Unit | 5.2 | 5 | 5 | Positive |
| **dC** | 228.1 | Wide | 112.1 | Unit | 1.7 | 5 | 5 | Positive |
| **[$^{13}C_9$]-dC** | 237.1 | Wide | 116.1 | Unit | 1.7 | 5 | 5 | Positive |
| **dA** | 252.1 | Wide | 136.1 | Unit | 6.7 | 12 | 5 | Positive |
| **[$^{15}N_5$-$^{13}C_{10}$]-dA** | 267.1 | Wide | 146.1 | Unit | 6.7 | 12 | 5 | Positive |
| **dG** | 268.0 | Wide | 152.1 | Unit | 5.0 | 6 | 5 | Positive |
| **[$^{15}N_5$-$^{13}C_{10}$]-dG** | 283.1 | Wide | 162.06 | Unit | 5.0 | 6 | 5 | Positive |
| **dT** | 243.1 | Wide | 127.1 | Unit | 5.6 | 3 | 5 | Positive |
| **[$^{15}N_2$-$^{13}C_{10}$]-dT** | 255.1 | Wide | 134.1 | Unit | 5.6 | 3 | 5 | Positive |

### switchSENSE® Chip surface functionalization

The binding kinetics experiments were performed on the heliX$^+$ instrument (Dynamic Biosensors GmbH, DE) using the switchSENSE® technology and a standard heliX® chip (ADP-48-2-0, Dynamic Biosensors), in which single-stranded DNA (anchor strands) are covalently attached to the chip surface. Each chip is equipped with 2 gold electrodes (or spots), with different DNA anchor strands. Herein, spot 1 was used as a measurement spot, with the dsDNA ligand strand elongated with the relevant Tet target sequence (dsDNA overhang containing 5mdC modification), and spot 2 as real-time reference spot, with unmodified dsDNA overhang, in order to monitor possible unspecific binding of Tet proteins (hpTet or DM T940A/Y1567F) on the anchor DNA and/or gold electrodes (Fig. S2 depicts the Adapter chip setup). The blue segment in spot 1 represents the modified DNA oligo hybridized to the DNA overhang of the ligand strand, whereas the pink dsDNA in spot 2 represents the unmodified DNA oligo hybridized to the DNA overhang of the ligand strand. For spot 1, the ligand strands elongated with **Tet target sequence** (5'– **CAT ACC GCA TTAC** TTT ATC AGT ACT TGT CAA CAC GAG CAG CCC GTA TAT TCT CCT ACA GCA CTA –3') were premixed with the modified complementary DNA overhang, whereas for spot 2 the ligand strands elongated with Tet target sequence were premixed with the unmodified complementary DNA overhang. The dsDNA ligand sequences with the overhangs were then mixed with the Adapter strand 1 and Adapter strand 2, carrying the red fluorophore, for hybridization to the spot 1 or 2, respectively (Dolcemascolo et al. 2023, and Higuera-Rodriguez et al., 2023). [7, 8] For complete hybridization, the DNA ligands were let at 25°C for at least 20 minutes on a shaker at 600 rpm. Secondly, the DNA ligands were immobilized on the biochip via hybridization of complementary anchor strand at a final concentration of 100 nM. The chip was regenerated and freshly functionalized before each measurement series. For chip regeneration, a high-pH regeneration solution (SOL-REG-1-5, Dynamic

Biosensors) was used to denature the double stranded DNA nanolevers, disrupting the hydrogen bonds between base pairs: the DNA ligands are washed away while the covalently attached single-stranded nanolevers remain on the surface and can be reused for a new functionalization step. Using FPS mode, a DNA-based biochip can be regenerated up to 50 times.

#### heliX$^+$ biosensor experiments and analysis

Interaction analysis was performed using a heliX$^+$ instrument (Dynamic Biosensors GmbH, DE) in fluorescence proximity sensing (FPS) mode (Dolcemascolo et al., 2023, Schulte et al., 2022). [7, 9] To repel the DNA ligands from the surface and ensure optimal accessibility for binding, a repulsive constant voltage of -0.2 V was applied.

In the FPS measurements, the Tet proteins (hpTet3 or DM T940A/Y1567F Tet3) were injected at increasing concentrations (hpTet3 at 62.5 nM – 125 nM – 250 nM, and DM T940A/Y1567F at 25 nM – 50 nM – 100 nM) after an initial blank (running buffer without protein) injection over the two electrodes of the biochip (Fig. S3 illustrates a typical measurement workflow). When the Tet proteins (hpTet3 or DM T940A/Y1567F Tet3) reach the modified DNA ligands, a decrease in fluorescence signal can be measured. The fluorescence signal changes occur on the timescale of seconds, and the time dependence of the fluorescence signal directly reflects the protein-DNA kinetics. Upon the injection of running buffer, the proteins dissociate, and a restoration of the original fluorescence signals can be observed.

For affinity studies, the sample tray containing the DNA ligands and protein samples was set to 15°C during measurements, while the experiment temperature on the biochip was set to 25°C. Protein analytes were diluted and measured in the relevant running buffer: kinetic studies were performed in HE140 (10 mM HEPES pH 7.4, 140 mM NaCl, 0.05 % Tween20, 50 µM EDTA, 50 µM EGTA); the salt dependency studies were performed in HE140 running buffer with increased NaCl concentration: HE200 (10 mM HEPES pH 7.4, 200 mM NaCl, 0.05 % Tween20, 50 µM EDTA, 50 µM EGTA), or HE300 (10 mM HEPES pH 7.4, 300 mM NaCl, 0.05 % Tween20, 50 µM EDTA, 50 µM EGTA). Flow rate for association and dissociation reactions was set to 200 µL/min. The red LED power was set to 2. Excitation and emission wavelengths for the red dye were 605-625 nm to 655-685 nm.

Experiment design, workflow and data analysis were performed with the heliOS software (Dynamic Biosensors GmbH, DE). The association and dissociation rates ($k_{on}$ and $k_{off}$), dissociation constants ($K_d$) and the respective error values were derived from a global exponential biphasic fit model, upon blank referencing correction.

3. SnapGene software (www.snapgene.com)

4. NEB Tm Calculator v1.16.7 (tmcalculator.neb.com)

5. Schröder, A.S., Steinbacher, J., Steigenberger, B., Gnerlich, F.A., Schiesser, S., Pfaffeneder, T., and Carell, T. (2014). Synthesis of a DNA promoter segment containing all four epigenetic nucleosides: 5-methyl-, 5-hydroxymethyl-, 5-formyl-, and 5-carboxy-2’-deoxycytidine. Angew. Chem. Int. Ed Engl. 53, 315–318.

6. Traube, F.R., Schiffers, S., Iwan, K., Kellner, S., Spada, F., Müller, M., and Carell, T. (2019). Isotope-dilution mass spectrometry for exact quantification of noncanonical DNA nucleosides. Nat. Protoc. 14, 283–312.

7. Dolcemascolo, R., Heras-Hernández, M., Goiriz, L., Montagud-Martínez, R., Requena-Menéndez, A., Ruiz, R., Pérez-Ràfols, A., Higuera-Rodríguez, R. A., Pérez-Ropero, G., Vranken, W. F., Martelli, T., Kaiser, W., Buijs, J., & Rodrigo, G. (2024). Repurposing the mammalian RNA-binding protein Musashi-1 as an allosteric translation repressor in bacteria. eLife, 12, RP91777.

8. Higuera-Rodriguez RA, De Pascali MC, Aziz M, Sattler M, Rant U, Kaiser W. Kinetic FRET Assay to Measure Binding-Induced Conformational Changes of Nucleic Acids. ACS Sens. 2023;8(12):4597-4606.

9. Schulte, C., Soldà, A., Spänig, S., Adams, N., Bekić, I., Streicher, W., Heider, D., Strasser, R., & Maric, H. M. (2022). Multivalent binding kinetics resolved by fluorescence proximity sensing. Communications biology, 5(1), 1070.
